# Quantifying the inverted U: A meta-analysis of prefrontal dopamine, D1-receptors, and working memory

**DOI:** 10.1101/2021.12.22.473899

**Authors:** Matthew A. Weber, Mackenzie M. Conlon, Hannah R. Stutt, Linder Wendt, Patrick Ten Eyck, Nandakumar S. Narayanan

## Abstract

Dopamine in the prefrontal cortex can be disrupted in human disorders that affect cognitive function such as Parkinson’s disease (PD), attention-deficit hyperactivity disorder (ADHD), and schizophrenia. Dopamine has a powerful effect on prefrontal circuits via the D1-type dopamine receptor (D1DR). It has been proposed that prefrontal dopamine has “inverted U-shaped” dynamics, with optimal dopamine and D1DR signaling required for optimal cognitive function. However, the quantitative relationship between prefrontal dopamine and cognitive function is not clear. Here, we conducted a meta-analysis of published manipulations of prefrontal dopamine and the effects on working memory, a high-level executive function in humans, primates, and rodents that involves maintaining and manipulating information over seconds to minutes. We reviewed 646 papers and found that 75 studies met criteria for inclusion. Our quantification of effect sizes for dopamine, D1DRs, and behavior revealed a negative quadratic slope. This is consistent with the proposed inverted U-shape of prefrontal dopamine and D1DRs and working memory performance, explaining 10% of the variance. Of note, the inverted quadratic fit was much stronger for prefrontal D1DRs alone, explaining 26% of the variance, compared to prefrontal dopamine alone, explaining 10% of the variance. Taken together, these data, derived from a variety of manipulations and systems, demonstrate that optimal prefrontal dopamine signalling is linked with higher cognitive function. Our results provide insight into the fundamental dynamics of prefrontal dopamine, which could be useful for pharmacological interventions targeting prefrontal dopaminergic circuits, and into the pathophysiology of human brain disease.

## Introduction

Human diseases that affect high-level cognitive processes such as working memory, reasoning, and flexibility can disrupt prefrontal dopamine. For instance, in humans with Parkinson’s disease, hypo- and hyperdopaminergic states have been linked with impaired cognition (Cools and D’Esposito, 2011; Mattay et al., 2002; Narayanan et al., 2013). In addition, dysfunctioning prefrontal dopaminergic systems may be related to the pathophysiology of attention-deficit hyperactivity disorder (ADHD) (Bellgrove et al., 2005), and prefrontal dopamine has been critically implicated in the pathogenesis of schizophrenia (Abi-Dargham et al., 2002; Goldman-Rakic et al., 2004; Okubo et al., 1997). Despite these data, the precise relationship between prefrontal dopamine and behavior is unclear. Understanding this relationship is relevant for pharmacological strategies that modulate prefrontal dopaminergic function to improve cognitive function in human disease (Soriano et al., 2010).

Preclinical work in rodents and non-human primates has established that prefrontal dopamine is required for high-level cognitive behaviors (Brozoski et al., 1979; Bubser and Schmidt, 1990; Kim et al., 2017). One of the most commonly studied cognitive behaviors is working memory, in which information is held for brief periods of time to guide future goal-directed behavior and has been studied extensively to show that decreased or increased prefrontal dopamine is linked with impaired behavioral performance (Cools and D’Esposito, 2011; Floresco, 2013; Goldman-Rakic et al., 2004). Prefrontal dopamine acts on cortical circuits via D1-type dopamine receptors (D1DRs), which also has been linked with impaired working memory performance (Floresco and Phillips, 2001; Goldman-Rakic et al., 2004; Seamans et al., 1998; Seamans and Yang, 2004). These findings lead to the hypothesis that working memory follows an inverted U-shaped function, in which optimal working memory performance is achieved with optimal levels of prefrontal dopamine and D1DR activation. While inverted U-shaped dynamics have substantial supporting evidence, the contours of this function are not clear. Further, it is not clear whether the inverted U-shape is more strongly dependent on either D1DR levels or overall prefrontal dopamine concentrations, or whether the curve is the same for both dopamine and D1DR manipulations. This is particularly relevant in predicting the degree of behavioral impairment that can be expected with prefrontal dopaminergic manipulations or for interventions that target D1DRs.

To formally quantify the relationship between prefrontal dopamine signaling and working memory, we conducted a meta-analysis of studies in which working memory and either prefrontal dopamine or D1DRs were measured. We report two major results: 1) there was a negative quadratic fit for the relationship between working memory and both prefrontal dopamine and prefrontal D1DR combined; and 2) the relationship was stronger for prefrontal D1DR manipulation and working memory, explaining 26% of the variance, compared to prefrontal dopamine and working memory that explained only 10% of the variance. We interpret these data in the context of prefrontal dopamine dynamics and their relevance for understanding prefrontal function in human disease.

## Methods

### Search strategy and inclusion/exclusion criteria

An electronic search of PubMed, PsychInfo, and Embase was performed on September 15, 2021 using the terms “frontal cortex,” “dopamine,” and “working memory”. Terms such as “human” and “dopamine D1” were also utilized to ensure a comprehensive search was completed. We restricted the search to peer-reviewed articles to ensure that only the most rigorous studies were included. Using functions in EndNote X9, we removed duplicates and literature reviews. resulting in 646 peer-reviewed articles. Two authors independently screened all of the abstracts (M.A.W and M.M.C) to determine appropriateness for this meta-analysis. We sought to synthesize data across multiple domains, including species of the model organism studied, working memory behavioral paradigms, and measure of prefrontal dopamine and D1DRs. Therefore, inclusion criteria were: 1) peer-reviewed original research in either rodents, non-human primates, or humans; that 2) measured prefrontal dopamine or D1DRs and 3) measured working-memory performance. Exclusion criteria were: 1) non-original research; 2) case studies; 3) in vitro or computational studies; 4) non-dopamine or D1DR studies; 5) studies that examined executive functions other than working memory; 6) studies that lacked between-group comparisons, control groups, or baseline measures; 7) central or peripheral pharmacology without direct measure of dopamine or D1DRs; and 8) study of genetic polymorphisms without direct measure of dopamine or D1DRs. This screening process resulted in 75 peer-reviewed publications included in the final quantitative analysis. This study’s design and hypothesis were not preregistered.

### Data extraction

Several variables were extracted from each study included in the final analysis. Broad characteristics of each study were: 1) article title; 2) authors; 3) publication year; 4) species; 5) experimental manipulation or comparison; 6) type of working memory task; and 7) type of prefrotnal dopamine or D1DR measure. Quantitative variables for the measure of working memory and prefrontal dopamine or D1DRs were: 1) number of subjects for each experimental group; 2) group average; and 3) group standard deviation or standard error. Every effort was taken to extract quantitative variables directly from the methods, results, and/or figure captions to ensure exact values were reported. Primary data extraction was completed by M.A.W, but all qualitative and quantitative data was verified independently by two other authors (H.R.S and N.S.N).

When multiple versions of the same working memory task were reported (e.g., the length of the working-memory delay period, see Abi-Dargham et al., 2002), we extracted the working memory behavior data points with the largest effect size. When multiple dopamine values were presented (e.g., at multiple time points during in vivo microdialysis, see Schmeichel et al., 2013), we extracted basal prefrontal dopamine values when available or data that matched the working memory time point as closely as possible when basal prefrontal levels were not reported. When the precise number of subjects in a group was not explicitly reported, we estimated group size based on the information available (e.g. Pietraszek et al., 2009). When group average, standard deviation, and standard error were not explicitly reported, we used plot digitizer software (Rohatgi, A., WebPlotDigitizer: Version 4.4, 2020, https://automeris.io/WebPlotDigitizer/) to extract relevant statistical data. Several publications contributed multiple data points to the final quantitative analysis because we were able to extract multiple values from these datasets. For example, Adams & Moghaddam, 1998, tested working memory performance at three time points following peripheral drug injection and included three corresponding prefrontal dopamine measures. Other examples include Novick et al., 2013 (two working memory paradigms), Szczepanik et al., 2020 (multiple doses of the same drug with corresponding prefrontal dopamine values), and Kellendonk et al., 2006 (multiple different measures of prefrontal dopamine - i.e., TH variscosities, D1 mRNA, DA content, *c-Fos* expression). We compared measures of working memory with prefrontal dopamine concentrations and D1DR activation in control and experimental groups, regardless of the specific statistical analysis that was presented in the publication. Our statistical analysis of control vs. experimental groups was used to generate effect sizes for both 1) difference in working memory performance and 2) difference in prefrontal dopamine or D1DRs between control and experimental conditions.

### Statistics

Following data extraction, we calculated Cohen’s D effect sizes for each measure of working memory and prefrontal dopamine or D1DRs. Effect sizes were adjusted so that enhanced working memory and increased prefrontal dopamine or D1DRs were reflected by positive values, and impaired working memory and dampened prefrontal dopamine or D1DRs were reflected by negative values. We then sorted effect sizes values based on prefrontal dopamine or D1DRs and grouped data to facilitate analysis of working memory performance (Tables 1 and 2).

**Table 1:** Quadratic equations (aX^2^ + bX + (Intercept)) derived from manipulations of prefrontal dopamine or D1DRs and measures of working memory performance.

|  | Coefficient | 95% Confidence Interval | p-value |
| --- | --- | --- | --- |
| Prefrontal D1DRs |  |  |  |
| a | -0.265 | -0.545, 0.015 | 0.063 |
| b | -0.353 | -0.539, -0.166 | <0.001 |
| (Intercept) | -0.513 | -1.097, 0.071 | 0.082 |
| Prefrontal Dopamine |  |  |  |
| a | 0.059 | -0.072, 0.191 | 0.374 |
| b | -0.091 | -0.146, -0.035 | 0.001 |
| (Intercept) | -0.843 | -1.117, -0.568 | <0.001 |
| Aggregate |  |  |  |
| a | -0.830 | -0.134, 0.089 | 0.69 |
| b | -0.120 | -0.171, -0.068 | <0.001 |
| (Intercept) | -0.830 | -1.080, -0.580 | <0.001 |

**Table 2:**
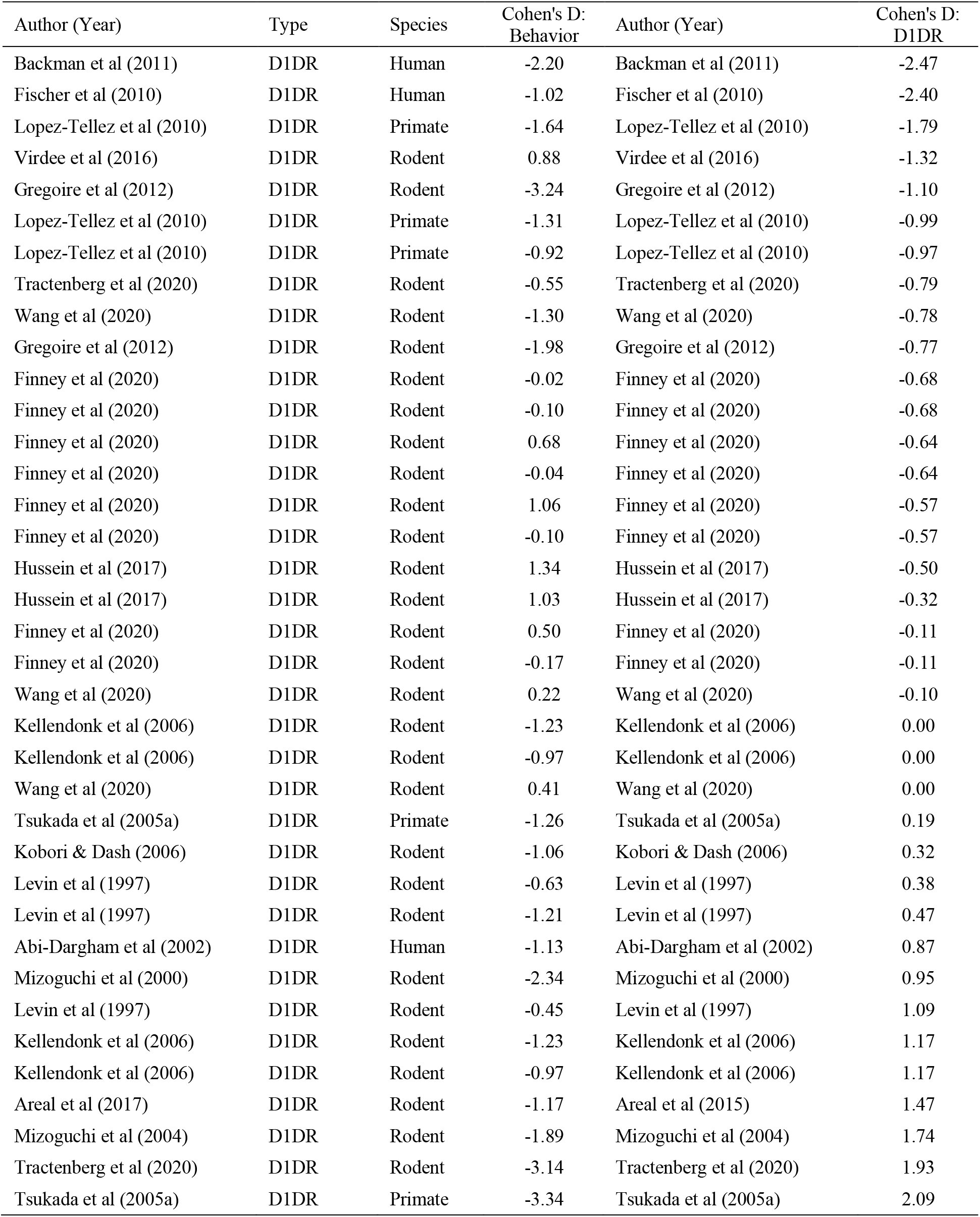
Studies that reported comparisons of prefrontal cortex D1-type dopamine receptors (D1DRs) and working memory between control and experimental subjects.

**Table 3:**
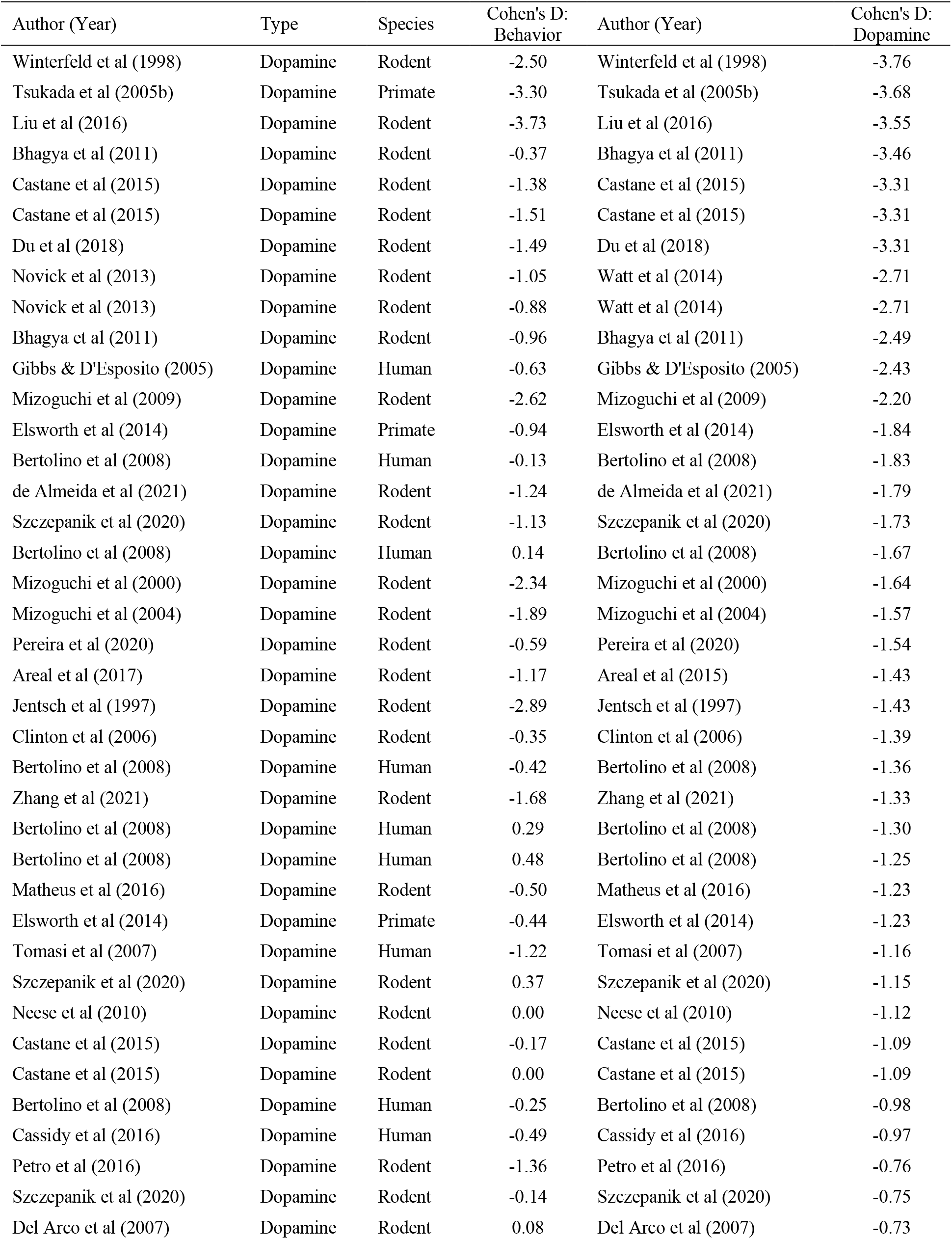
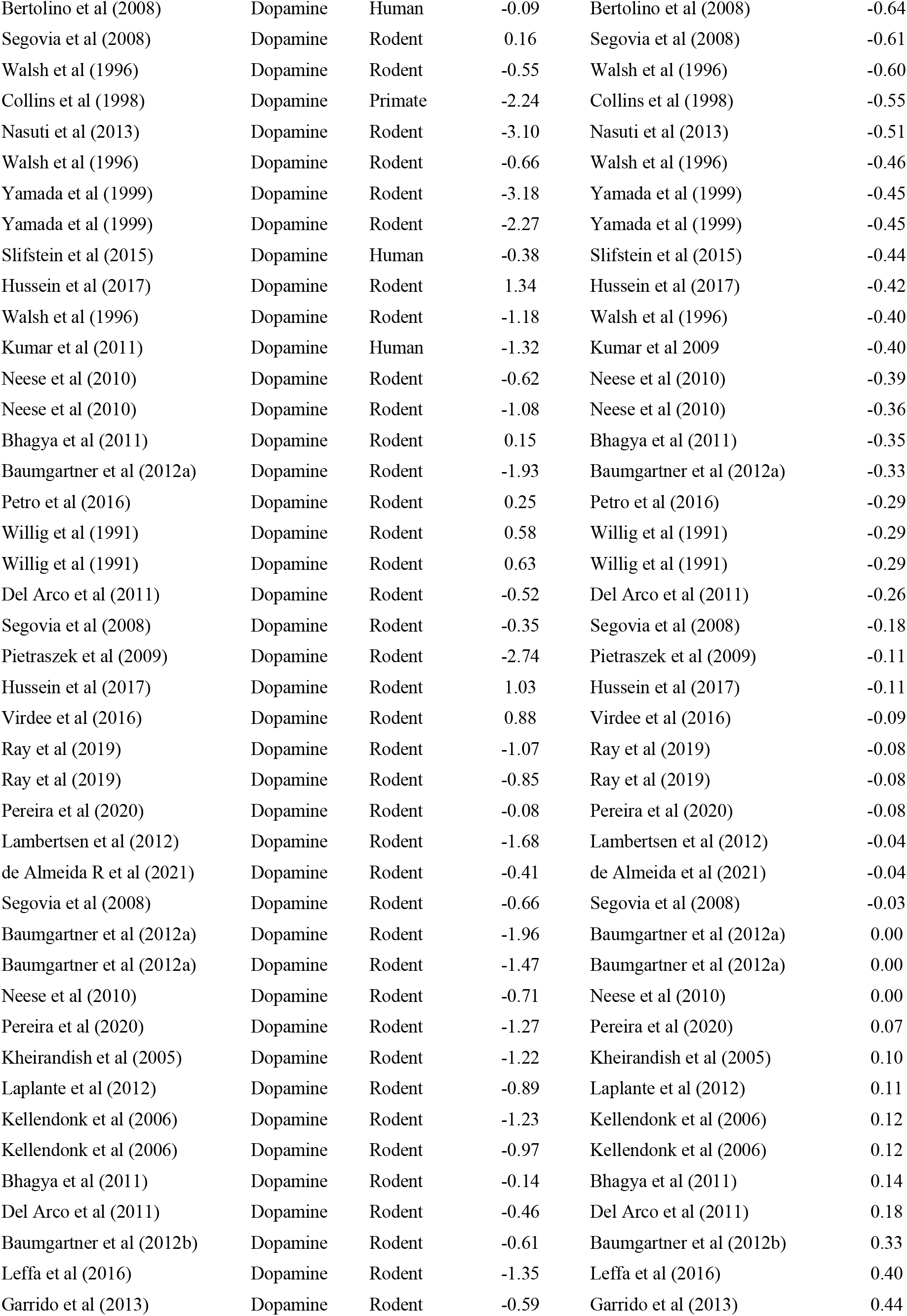
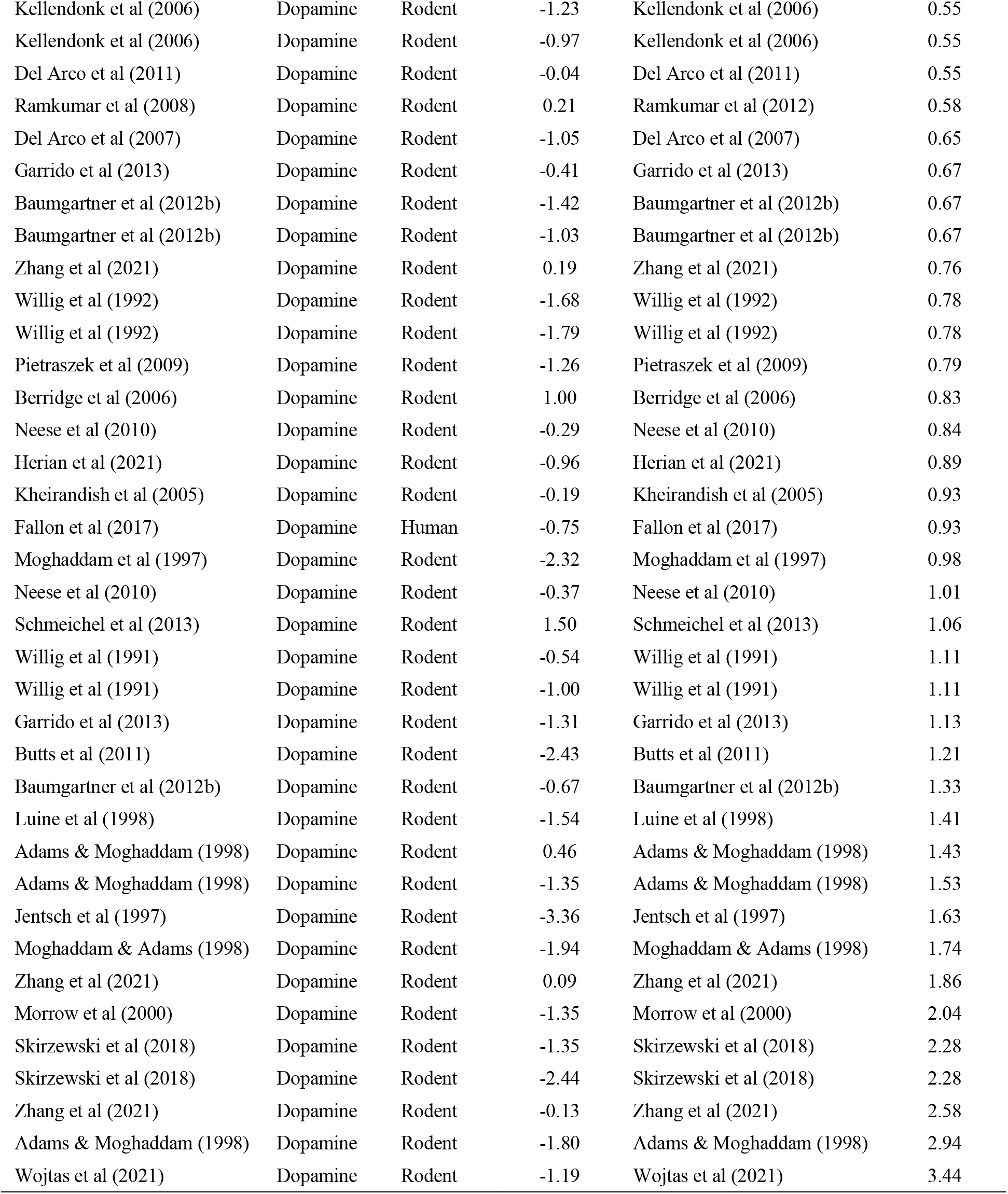
Studies that reported comparisons of prefrontal cortex dopamine and working memory between control and experimental subjects.

Statistical analyses were completed using R software, version 4.1.1. All code and raw data are available at https://narayanan.lab.uiowa.edu. All statistical analyses were performed and verified independently by the Biostatistics, Epidemiology, and Research Design Core within the Institute for Clinical and Translational Science at the University of Iowa.

The primary goal of this meta-analysis was to identify polynomial models (up to order three) that explain changes in working memory performance with changes in prefrontal dopamine and/or D1DRs. We developed models based on the relationship between working memory effect sizes and prefrontal dopamine and D1DR effect sizes. We excluded values greater than or less than a Cohen’s d of +/- 4, as these could have an outsized effect on our models. First, we fit a model based on working memory performance and all prefrontal dopamine and D1DRs. This analysis was followed by stratifying the data set to develop a model fit based on working memory performance and prefrontal dopamine and a model fit based on working memory performance and prefrontal D1DRs. Several publications contributed multiple values to the final data set, and this was accounted for by including a random intercept for each publication. Model fits were between different polynomial orders were compared via Akaike Information Criteria (AIC), with lower AICs indicating a better combination of parsimony and goodness of fit.

We used a bootstrap analysis approach to compare R^2^ values for prefrontal dopamine and prefrontal D1DRs. This process began by simulating a new dataset for both prefrontal dopamine and prefrontal D1DRs; we resampled the original datasets with replacement to create new datasets the same size as the original. Then, a quadratic model was built on each resampled dataset, and the R^2^ value of the dopamine model was subtracted from the R^2^ values of the D1DR model. This process was repeated 10,000 times to obtain bootstrap-estimated intervals that reflect 95% confidence for the difference between the two models and that one model’s fit is superior to the other. Here, a positive confidence interval that does not contain zero would indicate that the prefrontal D1DR model provides a superior R^2^ value compared to the prefrontal dopamine R^2^ value.

## Results

Our literature search and screening procedures yielded 75 journal articles that fit our criteria, resulting in 165 data points (Tables 1 and 2). After extreme values (Cohen’s d >+/- 4) were excluded, 156 data points remained. We found that a quadratic function provided the optimal model fit (2^nd^ order polynomial; p<0.001; AIC = 400.2 vs. linear AIC = 412.7). The R^2^ value for the negative quadratic fit was 0.10. A higher order polynomial model did not decrease AIC values (3^rd^ order AIC = 408.8), suggesting that the 2^nd^ order model is optimal.

We then stratified our data based on type of prefrontal measure, with a sub-analysis focused on prefrontal dopamine (i.e., dopamine content or turnover, tyrosine hydroxylase, dopamine transporter, etc.). These could include direct manipulations of prefrontal dopamine (e.g., dopamine depletion via 6-hydroxydopamine) or indirect manipulation such as stress or peripheral drug administration. For this analysis, we found 61 studies and 119 data points. A negative quadratic function provided the strongest fit with AIC = 314.4 (p<0.001; vs. linear AIC = 317.2, 3^rd^ order AIC = 322.7). The R^2^ value for our quadratic model was 0.10. No higher-order models yielded lower AIC values.

Prefrontal dopamine released from synaptic terminals can powerfully act on prefrontal D1DRs (Goldman-Rakic et al., 2004, p.; Seamans and Yang, 2004). We examined the role of prefrontal D1DR manipulations on working memory performance in 17 studies with 37 data points. In line with data on prefrontal dopamine, we found that a negative quadratic function again provided the best fit, with AIC = 102.6 (p<0.001; vs. linear AIC = 110.2; 3^rd^ order AIC = 106.3). The R^2^ value for this model was 0.26. Increasing the polynomial order coincided with an increase in the AIC values, suggesting that the negative quadratic model again provided the best combination of parsimony and goodness of fit. Adding an effect for the species being studied did not notably enhance our model’s goodness of fit, possibly due to insufficient sample size to detect this effect. When a variable controlling for species was added to our negative quadratic model, our AIC worsened from 314.4 to 315.5 for the prefrontal dopamine model and from 102.6 to 103.0 for the prefrontal D1DR model.

We then built new quadratic models using the resampling bootstrapped analysis described above for both prefrontal dopamine and prefrontal D1DRs and determined the difference between the two newly-built models. The average difference between R^2^ values for the 10,000 iterations was 0.14, where a positive value indicated that the prefrontal D1DR models had a greater R^2^ value. The 95% confidence interval for this result was (−0.10, 0.38) and the bootstrapped two-sided *p* value was 0.31.

## Discussion

Our goal was to quantify the relationship of working memory performance with prefrontal dopamine and D1DRs. We conducted a meta-analysis of 75 studies spanning rodents, non-human primates, and humans. These data suggest that 10% of the variance in working memory behavior was explained by manipulations of prefrontal dopamine, and 26% of the variance was explained by prefrontal D1DR manipulations. These data provide insight into how prefrontal dopamine and D1DRs affects cognitive behaviors.

These data are consistent with past work that has proposed inverted U-shaped relationship between prefrontal dopaminergic dynamics and working memory performance (Cools and D’Esposito, 2011; Floresco, 2013). We were able to demonstrate this idea by quantitatively fitting an inverted quadratic function, supporting the idea that there is an optimal regime for dopamine function in the prefrontal cortex that may facilitate a wide range of interacting synaptic and post-synaptic proteins (Arnsten et al., 2012; Arnsten and Li, 2005). In establishing this function, we show that prefrontal dopamine has strikingly different signaling principles than striatal dopamine (Kreitzer, 2009; Mohebi et al., 2019; Yahr et al., 1969), in which striatal dopamine depletion impairs movement and motivation, and increased dopamine facilitates movement and motivation.

While this work supports the hypothesis that working memory performance follows an inverted U-shape function dependent on prefrontal dopamine and D1DRs, our results should be interpreted carefully. For example, the bootstrapped analysis for models of prefrontal D1DRs were not significantly different from models of prefrontal dopamine; however, we note that there were fewer studies for prefrontal D1DRs, which may have affected our statistical power in separating prefrontal D1DRs from prefrontal dopamine manipulations. Another key constraint is that rodents do not have lateral prefrontal regions that are present in primates (Laubach et al., 2018), although dopamine is strongly released in medial prefrontal regions, and dopamine in these circuits may function according to similar principles (Floresco, 2013; Zahrt et al., 1997). It is also important to acknowledge that changes to working memory performance are not only impacted by manipulations of prefrontal dopamine and D1DRs. Other prefrontal dopamine receptors (Druzin et al., 2000; Glickstein et al., 2002), neurotransmitter systems (Monaco et al., 2015; Robbins and Arnsten, 2009), brain regions (Bolkan et al., 2017; Hart et al., 2018), and behaviors (i.e. interval timing, behavioral flexibility – Kim et al., 2017; Ragozzino, 2002; Zhang et al., 2019) are critical for optimal working memory performance. Furthermore, there are other paradigms that can be used to study other executive functions, and initial studies using set shifting, reversal learning, and interval timing imply that inverted U-shaped dynamics at least partially hold for these tasks, as well as others (Floresco, 2013; Parker et al., 2015; Robbins, 2007). However, our literature search revealed among manipulations of prefrontal dopamine and cognition, working memory paradigms had the largest number of studies, making it a reasonable starting point for comparisons across metholodogies and species. This work also has limitations that derive from comparing a broad range of studies across several different methodologies and model systems. However, this diversity is also a strength in that we report effects that are consistent across a range of approaches. Finally, publication bias may have affected this analysis, meaning that non-reviewed and unpublished research could have influenced our conclusions. While there are many small effect sizes within our datasets, the wealth of unpublished research possibly reporting nonsignificant prefrontal dopamine, prefrontal D1DR, or working memory changes could alter our interpretation of the inverted U-shape function.

In summary, this study advances the approach of bringing together diverse studies to elucidate patterns in prefrontal dopamine. A key finding here is that, while not statistically significant, the prefrontal D1DRs explained more variance than prefrontal dopamine. Fascinatingly, the initial description of the inverted-U shaped working memory function is based largely on pharmacological activation or inhibition of prefrontal D1DRs. It is possible that working memory performance is more strongly dependent on dopamine receptor activation than specific levels of prefrontal dopamine. This pattern will be useful in designing and interpreting preclinical studies, as well as in designing and optimizing new therapies for diseases such as ADHD, schizophrenia, and PD, which involve profound disruptions in prefrontal dopamine signaling.

## Acknowledgements and Author Note

MAW and NSN designed the meta-analysis. MAW and MMC independently screened abstracts for appropriateness. MAW collected the data, which was independently checked by NSN and HRS. LW, PTE, and NSN wrote the code and checked the analysis. MAW and NSN wrote the manuscript. HRS, LW, and PTE reviewed the manuscript. All code and raw data are available at https://narayanan.lab.uiowa.edu.

---

**Figure 1:**
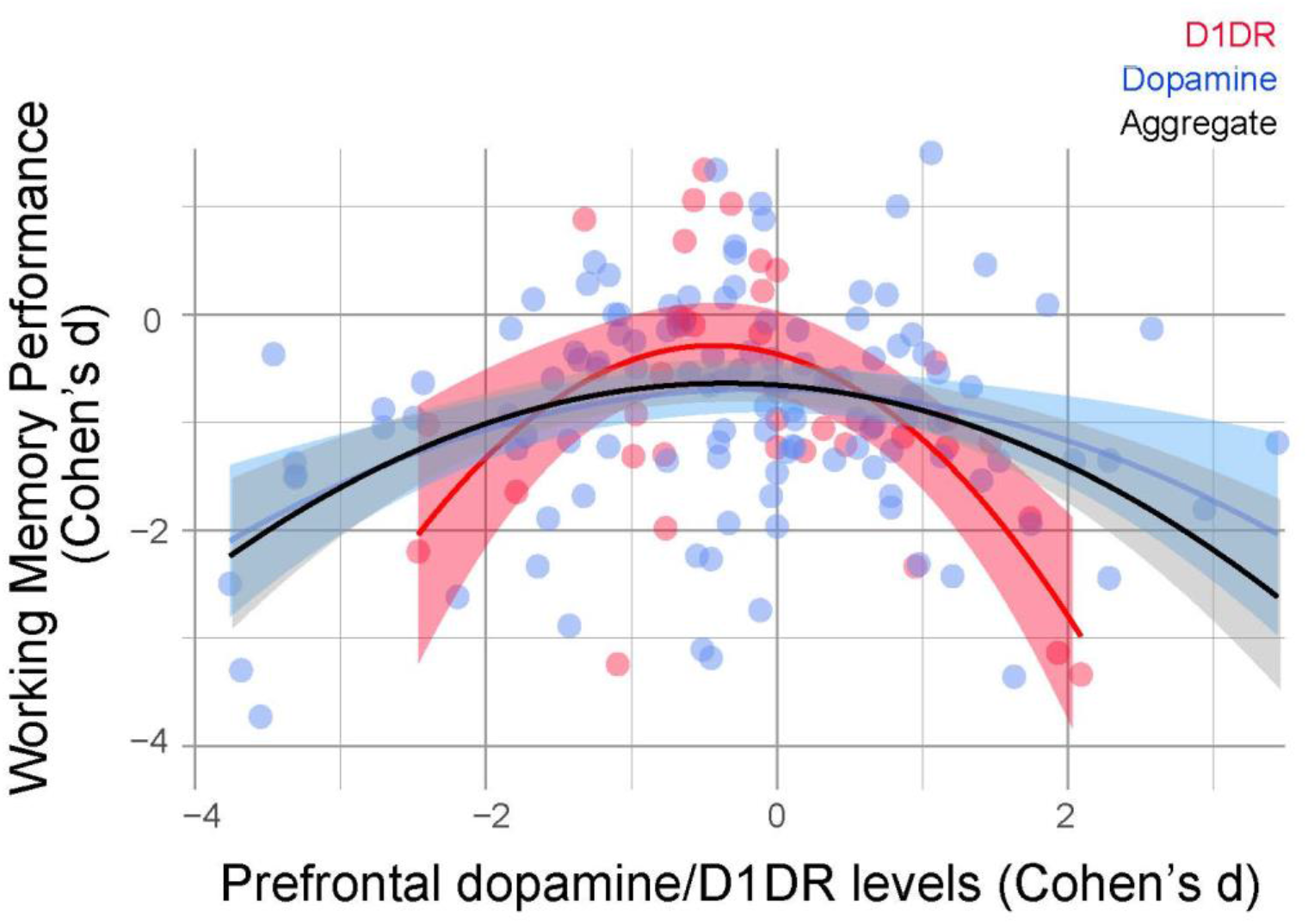
Working memory performance as a function of prefrontal dopamine and D1DRs. We included studies that measured both working memory performance and either prefrontal D1DRs or dopamine levels. We included studies from rodents, non-human primates, and humans, and expressed effect sizes in Cohen’s d. We found that studies that measured prefrontal D1DRs (red), prefrontal dopamine (blue) were best fit by a negative quadratic function. The model aggregating both prefrontal dopamine and D1DR measurements is shown in grey. Data from 75 studies and a total of 156 data points; 119 that measured prefrontal dopamine levels and 37 that measured prefrontal D1DR levels.

## References

Abi-Dargham, A., & Laruelle, M. (2005). Mechanisms of action of second generation antipsychotic drugs in schizophrenia: insights from brain imaging studies. Eur Psychiatry, 20(1), 15–27. doi:10.1016/j.eurpsy.2004.11.003

Abi-Dargham, A., Mawlawi, O., Lombardo, I., Gil, R., Martinez, D., Huang, Y., … Laruelle, M. (2002). Prefrontal dopamine D1 receptors and working memory in schizophrenia. J Neurosci, 22(9), 3708–3719. doi:10.1523/jneurosci.22-09-03708.2002

Adams, B., & Moghaddam, B. (1998). Corticolimbic dopamine neurotransmission is temporally dissociated from the cognitive and locomotor effects of phencyclidine. J Neurosci, 18(14), 5545–5554. doi:10.1523/jneurosci.18-14-05545.1998

Areal, L. B., Herlinger, A. L., Pelição, F. S., Martins-Silva, C., & Pires, R. G. W. (2017). Crack cocaine inhalation induces schizophrenia-like symptoms and molecular alterations in mice prefrontal cortex. J Psychiatr Res, 91, 57–63. doi:10.1016/j.jpsychires.2017.03.005

Arnsten, A. F., & Li, B. M. (2005). Neurobiology of executive functions: catecholamine influences on prefrontal cortical functions. Biol Psychiatry, 57(11), 1377–1384. doi:10.1016/j.biopsych.2004.08.019

Arnsten, A. F., Wang, M. J., & Paspalas, C. D. (2012). Neuromodulation of thought: flexibilities and vulnerabilities in prefrontal cortical network synapses. Neuron, 76(1), 223–239. doi:10.1016/j.neuron.2012.08.038

Bäckman, L., Karlsson, S., Fischer, H., Karlsson, P., Brehmer, Y., Rieckmann, A., … Nyberg, L. (2011). Dopamine D(1) receptors and age differences in brain activation during working memory. Neurobiol Aging, 32(10), 1849–1856. doi:10.1016/j.neurobiolaging.2009.10.018

Baumgartner, J., Smuts, C. M., Malan, L., Arnold, M., Yee, B. K., Bianco, L. E., … Zimmermann, M. B. (2012). Combined deficiency of iron and (n-3) fatty acids in male rats disrupts brain monoamine metabolism and produces greater memory deficits than iron deficiency or (n-3) fatty acid deficiency alone. J Nutr, 142(8), 1463–1471. doi:10.3945/jn.111.156281

Baumgartner, J., Smuts, C. M., Malan, L., Arnold, M., Yee, B. K., Bianco, L. E., … Zimmermann, M. B. (2012). In male rats with concurrent iron and (n-3) fatty acid deficiency, provision of either iron or (n-3) fatty acids alone alters monoamine metabolism and exacerbates the cognitive deficits associated with combined deficiency. J Nutr, 142(8), 1472–1478. doi:10.3945/jn.111.156299

Bellgrove, M. A., Hawi, Z., Kirley, A., Gill, M., & Robertson, I. H. (2005). Dissecting the attention deficit hyperactivity disorder (ADHD) phenotype: sustained attention, response variability and spatial attentional asymmetries in relation to dopamine transporter (DAT1) genotype. Neuropsychologia, 43(13), 1847–1857. doi:10.1016/j.neuropsychologia.2005.03.011

Berridge, C. W., Devilbiss, D. M., Andrzejewski, M. E., Arnsten, A. F., Kelley, A. E., Schmeichel, B., … Spencer, R. C. (2006). Methylphenidate preferentially increases catecholamine neurotransmission within the prefrontal cortex at low doses that enhance cognitive function. Biol Psychiatry, 60(10), 1111–1120. doi:10.1016/j.biopsych.2006.04.022

Bertolino, A., Di Giorgio, A., Blasi, G., Sambataro, F., Caforio, G., Sinibaldi, L., … Dallapiccola, B. (2008). Epistasis between dopamine regulating genes identifies a nonlinear response of the human hippocampus during memory tasks. Biol Psychiatry, 64(3), 226–234. doi:10.1016/j.biopsych.2008.02.001

Bhagya, V., Srikumar, B. N., Raju, T. R., & Shankaranarayana Rao, B. S. (2011). Chronic escitalopram treatment restores spatial learning, monoamine levels, and hippocampal long-term potentiation in an animal model of depression. Psychopharmacology, 214(2), 477–494. doi:10.1007/s00213-010-2054-x

Bolkan, S. S., Stujenske, J. M., Parnaudeau, S., Spellman, T. J., Rauffenbart, C., Abbas, A. I., … Kellendonk, C. (2017). Thalamic projections sustain prefrontal activity during working memory maintenance. Nat Neurosci, 20(7), 987–996. doi:10.1038/nn.4568

Brozoski, T. J., Brown, R. M., Rosvold, H. E., & Goldman, P. S. (1979). Cognitive deficit caused by regional depletion of dopamine in prefrontal cortex of rhesus monkey. Science, 205(4409), 929–932. doi:10.1126/science.112679

Bubser, M., & Schmidt, W. J. (1990). 6-Hydroxydopamine lesion of the rat prefrontal cortex increases locomotor activity, impairs acquisition of delayed alternation tasks, but does not affect uninterrupted tasks in the radial maze. Behav Brain Res, 37(2), 157–168. doi:10.1016/0166-4328(90)90091-r

Butts, K. A., Weinberg, J., Young, A. H., & Phillips, A. G. (2011). Glucocorticoid receptors in the prefrontal cortex regulate stress-evoked dopamine efflux and aspects of executive function. Proc Natl Acad Sci U S A, 108(45), 18459–18464. doi:10.1073/pnas.1111746108

Cassidy, C. M., Van Snellenberg, J. X., Benavides, C., Slifstein, M., Wang, Z., Moore, H., … Horga, G. (2016). Dynamic Connectivity between Brain Networks Supports Working Memory: Relationships to Dopamine Release and Schizophrenia. J Neurosci, 36(15), 4377–4388. doi:10.1523/jneurosci.3296-15.2016

Castañé, A., Santana, N., & Artigas, F. (2015). PCP-based mice models of schizophrenia: differential behavioral, neurochemical and cellular effects of acute and subchronic treatments. Psychopharmacology (Berl), 232(21-22), 4085–4097. doi:10.1007/s00213-015-3946-6

Castner, S. A., Vosler, P. S., & Goldman-Rakic, P. S. (2005). Amphetamine sensitization impairs cognition and reduces dopamine turnover in primate prefrontal cortex. Biol Psychiatry, 57(7), 743–751. doi:10.1016/j.biopsych.2004.12.019

Clinton, S. M., Sucharski, I. L., & Finlay, J. M. (2006). Desipramine attenuates working memory impairments induced by partial loss of catecholamines in the rat medial prefrontal cortex. Psychopharmacology (Berl), 183(4), 404–412. doi:10.1007/s00213-005-0221-2

Collins, P., Roberts, A. C., Dias, R., Everitt, B. J., & Robbins, T. W. (1998). Perseveration and strategy in a novel spatial self-ordered sequencing task for nonhuman primates: effects of excitotoxic lesions and dopamine depletions of the prefrontal cortex. J Cogn Neurosci, 10(3), 332–354. doi:10.1162/089892998562771

Cools, R., & D’Esposito, M. (2011). Inverted-U-shaped dopamine actions on human working memory and cognitive control. Biol Psychiatry, 69(12), e113–125. doi:10.1016/j.biopsych.2011.03.028

de Almeida, G. R. L., Szczepanik, J. C., Selhorst, I., Schmitz, A. E., Dos Santos, B., Cunha, M. P., … Dafre, A. L. (2021). Methylglyoxal-Mediated Dopamine Depletion, Working Memory Deficit, and Depression-Like Behavior Are Prevented by a Dopamine/Noradrenaline Reuptake Inhibitor. Mol Neurobiol, 58(2), 735–749. doi:10.1007/s12035-020-02146-3

Del Arco, A., Segovia, G., de Blas, M., Garrido, P., Acuña-Castroviejo, D., Pamplona, R., & Mora, F. (2011). Prefrontal cortex, caloric restriction and stress during aging: studies on dopamine and acetylcholine release, BDNF and working memory. Behav Brain Res, 216(1), 136–145. doi:10.1016/j.bbr.2010.07.024

Del Arco, A., Segovia, G., Garrido, P., de Blas, M., & Mora, F. (2007). Stress, prefrontal cortex and environmental enrichment: studies on dopamine and acetylcholine release and working memory performance in rats. Behav Brain Res, 176(2), 267–273. doi:10.1016/j.bbr.2006.10.006

Druzin, M. Y., Kurzina, N. P., Malinina, E. P., & Kozlov, A. P. (2000). The effects of local application of D2 selective dopaminergic drugs into the medial prefrontal cortex of rats in a delayed spatial choice task. Behav Brain Res, 109(1), 99–111. doi:10.1016/s0166-4328(99)00166-7

Du, C. X., Liu, J., Guo, Y., Zhang, L., & Zhang, Q. J. (2018). Lesions of the lateral habenula improve working memory performance in hemiparkinsonian rats. Neurosci Lett, 662, 162–166. doi:10.1016/j.neulet.2017.10.027

Elsworth, J. D., Groman, S. M., Jentsch, J. D., Leranth, C., Redmond, D. E., Jr., Kim, J. D., … Roth, R. H. (2014). Primate phencyclidine model of schizophrenia: sex-specific effects on cognition, brain derived neurotrophic factor, spine synapses, and dopamine turnover in prefrontal cortex. Int J Neuropsychopharmacol, 18(6). doi:10.1093/ijnp/pyu048

Fallon, S. J., van der Schaaf, M. E., Ter Huurne, N., & Cools, R. (2017). The Neurocognitive Cost of Enhancing Cognition with Methylphenidate: Improved Distractor Resistance but Impaired Updating. J Cogn Neurosci, 29(4), 652–663. doi:10.1162/jocn_a_01065

Finney, C. A., Proschogo, N. W., Snepvangers, A., Holmes, N. M., Westbrook, R. F., & Clemens, K. J. (2020). The effect of standard laboratory diets on estrogen signaling and spatial memory in male and female rats. Physiol Behav, 215, 112787. doi:10.1016/j.physbeh.2019.112787

Fischer, H., Nyberg, L., Karlsson, S., Karlsson, P., Brehmer, Y., Rieckmann, A., … Bäckman, L. (2010). Simulating Neurocognitive Aging: Effects of a Dopaminergic Antagonist on Brain Activity During Working Memory. Biological Psychiatry, 67(6), 575–580. doi:10.1016/j.biopsych.2009.12.013

Floresco, S. B. (2013). Prefrontal dopamine and behavioral flexibility: shifting from an “inverted-U” toward a family of functions. Front Neurosci, 7, 62. doi:10.3389/fnins.2013.00062

Floresco, S. B., & Phillips, A. G. (2001). Delay-dependent modulation of memory retrieval by infusion of a dopamine D1 agonist into the rat medial prefrontal cortex. Behav Neurosci, 115(4), 934–939. Retrieved from https://www.ncbi.nlm.nih.gov/pubmed/11508732

Garrido, P., De Blas, M., Ronzoni, G., Cordero, I., Antón, M., Giné, E., … Mora, F. (2013). Differential effects of environmental enrichment and isolation housing on the hormonal and neurochemical responses to stress in the prefrontal cortex of the adult rat: relationship to working and emotional memories. J Neural Transm (Vienna), 120(5), 829–843. doi:10.1007/s00702-012-0935-3

Gibbs, S. E., & D’Esposito, M. (2005). Individual capacity differences predict working memory performance and prefrontal activity following dopamine receptor stimulation. Cogn Affect Behav Neurosci, 5(2), 212–221. doi:10.3758/cabn.5.2.212

Glickstein, S. B., Hof, P. R., & Schmauss, C. (2002). Mice lacking dopamine D2 and D3 receptors have spatial working memory deficits. J Neurosci, 22(13), 5619–5629. doi:20026543

Goldman-Rakic, P. S., Castner, S. A., Svensson, T. H., Siever, L. J., & Williams, G. V. (2004). Targeting the dopamine D1 receptor in schizophrenia: insights for cognitive dysfunction. Psychopharmacology (Berl), 174(1), 3–16. doi:10.1007/s00213-004-1793-y

Grégoire, S., Rivalan, M., Le Moine, C., & Dellu-Hagedorn, F. (2012). The synergy of working memory and inhibitory control: behavioral, pharmacological and neural functional evidences. Neurobiol Learn Mem, 97(2), 202–212. doi:10.1016/j.nlm.2011.12.003

Hart, G., Bradfield, L. A., & Balleine, B. W. (2018). Prefrontal Corticostriatal Disconnection Blocks the Acquisition of Goal-Directed Action. J Neurosci, 38(5), 1311–1322. doi:10.1523/JNEUROSCI.2850-17.2017

Herian, M., Skawski, M., Wojtas, A., Sobocińska, M. K., Noworyta, K., & Gołembiowska, K. (2021). Tolerance to neurochemical and behavioral effects of the hallucinogen 25I-NBOMe. Psychopharmacology (Berl), 238(8), 2349–2364. doi:10.1007/s00213-021-05860-5

Hussein, A. M., Aher, Y. D., Kalaba, P., Aher, N. Y., Dragačević, V., Radoman, B., … Lubec, G. (2017). A novel heterocyclic compound improves working memory in the radial arm maze and modulates the dopamine receptor D1R in frontal cortex of the Sprague-Dawley rat. Behav Brain Res, 332, 308–315. doi:10.1016/j.bbr.2017.06.023

Jentsch, J. D., Andrusiak, E., Tran, A., Bowers, M. B., Jr., & Roth, R. H. (1997). Delta 9-tetrahydrocannabinol increases prefrontal cortical catecholaminergic utilization and impairs spatial working memory in the rat: blockade of dopaminergic effects with HA966. Neuropsychopharmacology, 16(6), 426–432. doi:10.1016/s0893-133x(97)00018-3

Kadowaki Horita, T., Kobayashi, M., Mori, A., Jenner, P., & Kanda, T. (2013). Effects of the adenosine A2A antagonist istradefylline on cognitive performance in rats with a 6-OHDA lesion in prefrontal cortex. Psychopharmacology (Berl), 230(3), 345–352. doi:10.1007/s00213-013-3158-x

Kellendonk, C., Simpson, E. H., Polan, H. J., Malleret, G., Vronskaya, S., Winiger, V., … Kandel, E. R. (2006). Transient and selective overexpression of dopamine D2 receptors in the striatum causes persistent abnormalities in prefrontal cortex functioning. Neuron, 49(4), 603–615. doi:10.1016/j.neuron.2006.01.023

Kheirandish, L., Gozal, D., Pequignot, J. M., Pequignot, J., & Row, B. W. (2005). Intermittent hypoxia during development induces long-term alterations in spatial working memory, monoamines, and dendritic branching in rat frontal cortex. Pediatr Res, 58(3), 594–599. doi:10.1203/01.pdr.0000176915.19287.e2

Kim, Y. C., Han, S. W., Alberico, S. L., Ruggiero, R. N., De Corte, B., Chen, K. H., & Narayanan, N. S. (2017). Optogenetic Stimulation of Frontal D1 Neurons Compensates for Impaired Temporal Control of Action in Dopamine-Depleted Mice. Curr Biol, 27(1), 39–47. doi:10.1016/j.cub.2016.11.029

Kobori, N., & Dash, P. K. (2006). Reversal of brain injury-induced prefrontal glutamic acid decarboxylase expression and working memory deficits by D1 receptor antagonism. J Neurosci, 26(16), 4236–4246. doi:10.1523/jneurosci.4687-05.2006

Kreitzer, A. C. (2009). Physiology and pharmacology of striatal neurons. Annu Rev Neurosci, 32, 127–147. doi:10.1146/annurev.neuro.051508.135422

Kumar, A. M., Fernandez, J. B., Singer, E. J., Commins, D., Waldrop-Valverde, D., Ownby, R. L., & Kumar, M. (2009). Human immunodeficiency virus type 1 in the central nervous system leads to decreased dopamine in different regions of postmortem human brains. J Neurovirol, 15(3), 257–274. doi:10.1080/13550280902973952

Kumar, A. M., Ownby, R. L., Waldrop-Valverde, D., Fernandez, B., & Kumar, M. (2011). Human immunodeficiency virus infection in the CNS and decreased dopamine availability: relationship with neuropsychological performance. J Neurovirol, 17(1), 26–40. doi:10.1007/s13365-010-0003-4

Lambertsen, K. L., Gramsbergen, J. B., Sivasaravanaparan, M., Ditzel, N., Sevelsted-Møller, L. M., Oliván-Viguera, A., … Köhler, R. (2012). Genetic KCa3.1-deficiency produces locomotor hyperactivity and alterations in cerebral monoamine levels. PLoS One, 7(10), e47744. doi:10.1371/journal.pone.0047744

Laplante, F., Zhang, Z. W., Huppé-Gourgues, F., Dufresne, M. M., Vaucher, E., & Sullivan, R. M. (2012). Cholinergic depletion in nucleus accumbens impairs mesocortical dopamine activation and cognitive function in rats. Neuropharmacology, 63(6), 1075–1084. doi:10.1016/j.neuropharm.2012.07.033

Laubach, M., Amarante, L. M., Swanson, K., & White, S. R. (2018). What, If Anything, Is Rodent Prefrontal Cortex? eNeuro, 5(5). doi:10.1523/ENEURO.0315-18.2018

Leffa, D. T., de Souza, A., Scarabelot, V. L., Medeiros, L. F., de Oliveira, C., Grevet, E. H., … Torres, I. L. S. (2016). Transcranial direct current stimulation improves short-term memory in an animal model of attention-deficit/hyperactivity disorder. Eur Neuropsychopharmacol, 26(2), 368–377. doi:10.1016/j.euroneuro.2015.11.012

Levin, E. D., Torry, D., Christopher, N. C., Yu, X., Einstein, G., & Schwartz-Bloom, R. D. (1997). Is binding to nicotinic acetylcholine and dopamine receptors related to working memory in rats? Brain Res Bull, 43(3), 295–304. doi:10.1016/s0361-9230(97)00009-9

Liu, K. C., Li, J. Y., Xie, W., Li, L. B., Zhang, J., Du, C. X., … Zhang, L. (2016). Activation and blockade of serotonin(6) receptors in the dorsal hippocampus enhance T maze and hole-board performance in a unilateral 6-hydroxydopamine rat model of Parkinson’s disease. Brain Res, 1650, 184–195. doi:10.1016/j.brainres.2016.09.009

Liu, Y., Liu, J., Jiao, S. R., Liu, X., Guo, Y., Zhang, J., … Zhang, L. (2019). Serotonin(1A) receptors in the dorsal hippocampus regulate working memory and long-term habituation in the hemiparkinsonian rats. Behav Brain Res, 376, 112207. doi:10.1016/j.bbr.2019.112207

López-Téllez, J. F., López-Aranda, M. F., Navarro-Lobato, I., Masmudi-Martín, M., Montañez, E. M., Calvo, E. B., & Khan, Z. U. (2010). Prefrontal inositol triphosphate is molecular correlate of working memory in nonhuman primates. J Neurosci, 30(8), 3067–3071. doi:10.1523/jneurosci.4565-09.2010

Luine, V. N., Richards, S. T., Wu, V. Y., & Beck, K. D. (1998). Estradiol enhances learning and memory in a spatial memory task and effects levels of monoaminergic neurotransmitters. Hormones and Behavior, 34(2), 149–162. doi:10.1006/hbeh.1998.1473

Matheus, F. C., Rial, D., Real, J. I., Lemos, C., Ben, J., Guaita, G. O., … Prediger, R. D. (2016). Decreased synaptic plasticity in the medial prefrontal cortex underlies short-term memory deficits in 6-OHDA-lesioned rats. Behav Brain Res, 301, 43–54. doi:10.1016/j.bbr.2015.12.011

Mattay, V. S., Tessitore, A., Callicott, J. H., Bertolino, A., Goldberg, T. E., Chase, T. N., … Weinberger, D. R. (2002). Dopaminergic modulation of cortical function in patients with Parkinson’s disease. Ann Neurol, 51(2), 156–164. doi:10.1002/ana.10078

Milstein, J. A., Elnabawi, A., Vinish, M., Swanson, T., Enos, J. K., Bailey, A. M., … Frost, D. O. (2013). Olanzapine treatment of adolescent rats causes enduring specific memory impairments and alters cortical development and function. PLoS One, 8(2), e57308. doi:10.1371/journal.pone.0057308

Mizoguchi, K., Ishige, A., Takeda, S., Aburada, M., & Tabira, T. (2004). Endogenous glucocorticoids are essential for maintaining prefrontal cortical cognitive function. J Neurosci, 24(24), 5492–5499. doi:10.1523/jneurosci.0086-04.2004

Mizoguchi, K., Shoji, H., Tanaka, Y., Maruyama, W., & Tabira, T. (2009). Age-related spatial working memory impairment is caused by prefrontal cortical dopaminergic dysfunction in rats. Neuroscience, 162(4), 1192–1201. doi:10.1016/j.neuroscience.2009.05.023

Mizoguchi, K., Yuzurihara, M., Ishige, A., Sasaki, H., Chui, D. H., & Tabira, T. (2000). Chronic stress induces impairment of spatial working memory because of prefrontal dopaminergic dysfunction. J Neurosci, 20(4), 1568–1574. doi:10.1523/jneurosci.20-04-01568.2000

Moghaddam, B., Adams, B., Verma, A., & Daly, D. (1997). Activation of glutamatergic neurotransmission by ketamine: A novel step in the pathway from NMDA receptor blockade to dopaminergic and cognitive disruptions associated with the prefrontal cortex. Journal of Neuroscience, 17(8), 2921–2927. doi:10.1523/jneurosci.17-08-02921.1997

Moghaddam, B., & Adams, B. W. (1998). Reversal of phencyclidine effects by a group II metabotropic glutamate receptor agonist in rats. Science, 281(5381), 1349–1352. doi:10.1126/science.281.5381.1349

Mohebi, A., Pettibone, J. R., Hamid, A. A., Wong, J. T., Vinson, L. T., Patriarchi, T., … Berke, J. D. (2019). Dissociable dopamine dynamics for learning and motivation. Nature, 570(7759), 65–70. doi:10.1038/s41586-019-1235-y

Monaco, S. A., Gulchina, Y., & Gao, W. J. (2015). NR2B subunit in the prefrontal cortex: A double-edged sword for working memory function and psychiatric disorders. Neurosci Biobehav Rev, 56, 127–138. doi:10.1016/j.neubiorev.2015.06.022

Morrow, B. A., Roth, R. H., & Elsworth, J. D. (2000). TMT, a predator odor, elevates mesoprefrontal dopamine metabolic activity and disrupts short-term working memory in the rat. Brain Res Bull, 52(6), 519–523. doi:10.1016/s0361-9230(00)00290-2

Narayanan, N. S., Rodnitzky, R. L., & Uc, E. Y. (2013). Prefrontal dopamine signaling and cognitive symptoms of Parkinson’s disease. Rev Neurosci, 24(3), 267–278. doi:10.1515/revneuro-2013-0004

Nasuti, C., Carloni, M., Fedeli, D., Gabbianelli, R., Di Stefano, A., Serafina, C. L., … Ciccocioppo, R. (2013). Effects of early life permethrin exposure on spatial working memory and on monoamine levels in different brain areas of pre-senescent rats. Toxicology, 303, 162–168. doi:10.1016/j.tox.2012.09.016

Neese, S. L., Wang, V. C., Doerge, D. R., Woodling, K. A., Andrade, J. E., Helferich, W. G., … Schantz, S. L. (2010). Impact of dietary genistein and aging on executive function in rats. Neurotoxicol Teratol, 32(2), 200–211. doi:10.1016/j.ntt.2009.11.003

Novick, A. M., Miiller, L. C., Forster, G. L., & Watt, M. J. (2013). Adolescent social defeat decreases spatial working memory performance in adulthood. Behav Brain Funct, 9, 39. doi:10.1186/1744-9081-9-39

Okubo, Y., Suhara, T., Suzuki, K., Kobayashi, K., Inoue, O., Terasaki, O., … Toru, M. (1997). Decreased prefrontal dopamine D1 receptors in schizophrenia revealed by PET. Nature, 385(6617), 634–636. doi:10.1038/385634a0

Parker, K. L., Ruggiero, R. N., & Narayanan, N. S. (2015). Infusion of D1 Dopamine Receptor Agonist into Medial Frontal Cortex Disrupts Neural Correlates of Interval Timing. Front Behav Neurosci, 9, 294. doi:10.3389/fnbeh.2015.00294

Pereira, A. G., Poli, A., Matheus, F. C., de Bortoli da Silva, L., Fadanni, G. P., Izídio, G. S., … Prediger, R. D. (2020). Temporal development of neurochemical and cognitive impairments following reserpine administration in rats. Behav Brain Res, 383, 112517. doi:10.1016/j.bbr.2020.112517

Petro, A., Sexton, H. G., Miranda, C., Rastogi, A., Freedman, J. H., & Levin, E. D. (2016). Persisting neurobehavioral effects of developmental copper exposure in wildtype and metallothionein 1 and 2 knockout mice. BMC Pharmacol Toxicol, 17(1), 55. doi:10.1186/s40360-016-0096-3

Pietraszek, M., Michaluk, J., Romańska, I., Wasik, A., Gołembiowska, K., & Antkiewicz-Michaluk, L. (2009). 1-Methyl-1,2,3,4-tetrahydroisoquinoline antagonizes a rise in brain dopamine metabolism, glutamate release in frontal cortex and locomotor hyperactivity produced by MK-801 but not the disruptions of prepulse inhibition, and impairment of working memory in rat. Neurotox Res, 16(4), 390–407. doi:10.1007/s12640-009-9097-y

R Core Team (2021). R: A language and environment for statistical computing. R Foundation for Statistical Computing, Vienna, Austria. https://www.R-project.org/

Ragozzino, M. E. (2002). The effects of dopamine D(1) receptor blockade in the prelimbic-infralimbic areas on behavioral flexibility. Learn Mem, 9(1), 18–28. doi:10.1101/lm.45802

Ramkumar, K., Srikumar, B. N., Shankaranarayana Rao, B. S., & Raju, T. R. (2008). Self-stimulation rewarding experience restores stress-induced CA3 dendritic atrophy, spatial memory deficits and alterations in the levels of neurotransmitters in the hippocampus. Neurochem Res, 33(9), 1651–1662. doi:10.1007/s11064-007-9511-x

Ramkumar, K., Srikumar, B. N., Venkatasubramanian, D., Siva, R., Shankaranarayana Rao, B. S., & Raju, T. R. (2012). Reversal of stress-induced dendritic atrophy in the prefrontal cortex by intracranial self-stimulation. J Neural Transm (Vienna), 119(5), 533–543. doi:10.1007/s00702-011-0740-4

Ray, A., Chitre, N. M., Daphney, C. M., Blough, B. E., Canal, C. E., & Murnane, K. S. (2019). Effects of the second-generation “bath salt” cathinone alpha-pyrrolidinopropiophenone (α-PPP) on behavior and monoamine neurochemistry in male mice. Psychopharmacology (Berl), 236(3), 1107–1117. doi:10.1007/s00213-018-5044-z

Robbins, T. W., & Arnsten, A. F. (2009). The neuropsychopharmacology of fronto-executive function: monoaminergic modulation. Annu Rev Neurosci, 32, 267–287. doi:10.1146/annurev.neuro.051508.135535

Robbins, T. W., & Roberts, A. C. (2007). Differential regulation of fronto-executive function by the monoamines and acetylcholine. Cereb Cortex, 17 Suppl 1, i151–160. doi:10.1093/cercor/bhm066

Schmeichel, B. E., Zemlan, F. P., & Berridge, C. W. (2013). A selective dopamine reuptake inhibitor improves prefrontal cortex-dependent cognitive function: potential relevance to attention deficit hyperactivity disorder. Neuropharmacology, 64(1), 321–328. doi:10.1016/j.neuropharm.2012.07.005

Seamans, J. K., Floresco, S. B., & Phillips, A. G. (1998). D1 receptor modulation of hippocampal-prefrontal cortical circuits integrating spatial memory with executive functions in the rat. J Neurosci, 18(4), 1613–1621. Retrieved from https://www.ncbi.nlm.nih.gov/pubmed/9454866

Seamans, J. K., & Yang, C. R. (2004). The principal features and mechanisms of dopamine modulation in the prefrontal cortex. Prog Neurobiol, 74(1), 1–58. doi:10.1016/j.pneurobio.2004.05.006

Segovia, G., Del Arco, A., de Blas, M., Garrido, P., & Mora, F. (2008). Effects of an enriched environment on the release of dopamine in the prefrontal cortex produced by stress and on working memory during aging in the awake rat. Behav Brain Res, 187(2), 304–311. doi:10.1016/j.bbr.2007.09.024

Skirzewski, M., Karavanova, I., Shamir, A., Erben, L., Garcia-Olivares, J., Shin, J. H., … Buonanno, A. (2018). ErbB4 signaling in dopaminergic axonal projections increases extracellular dopamine levels and regulates spatial/working memory behaviors. Mol Psychiatry, 23(11), 2227–2237. doi:10.1038/mp.2017.132

Slifstein, M., van de Giessen, E., Van Snellenberg, J., Thompson, J. L., Narendran, R., Gil, R., … Abi-Dargham, A. (2015). Deficits in prefrontal cortical and extrastriatal dopamine release in schizophrenia: a positron emission tomographic functional magnetic resonance imaging study. JAMA Psychiatry, 72(4), 316–324. doi:10.1001/jamapsychiatry.2014.2414

Soriano, A., Vendrell, M., Gonzalez, S., Mallol, J., Albericio, F., Royo, M., … Casado, V. (2010). A hybrid indoloquinolizidine peptide as allosteric modulator of dopamine D1 receptors. J Pharmacol Exp Ther, 332(3), 876–885. doi:10.1124/jpet.109.158824

Szczepanik, J. C., de Almeida, G. R. L., Cunha, M. P., & Dafre, A. L. (2020). Repeated Methylglyoxal Treatment Depletes Dopamine in the Prefrontal Cortex, and Causes Memory Impairment and Depressive-Like Behavior in Mice. Neurochem Res, 45(2), 354–370. doi:10.1007/s11064-019-02921-2

Tomasi, D., Goldstein, R. Z., Telang, F., Maloney, T., Alia-Klein, N., Caparelli, E. C., & Volkow, N. D. (2007). Widespread disruption in brain activation patterns to a working memory task during cocaine abstinence. Brain Res, 1171, 83–92. doi:10.1016/j.brainres.2007.06.102

Tomasi, D., Volkow, N. D., Wang, G. J., Wang, R., Telang, F., Caparelli, E. C., … Fowler, J. S. (2011). Methylphenidate enhances brain activation and deactivation responses to visual attention and working memory tasks in healthy controls. Neuroimage, 54(4), 3101–3110. doi:10.1016/j.neuroimage.2010.10.060

Tractenberg, S. G., Orso, R., Creutzberg, K. C., Malcon, L. M. C., Lumertz, F. S., Wearick-Silva, L. E., … Grassi-Oliveira, R. (2020). Vulnerable and resilient cognitive performance related to early life stress: The potential mediating role of dopaminergic receptors in the medial prefrontal cortex of adult mice. Int J Dev Neurosci, 80(1), 13–27. doi:10.1002/jdn.10004

Tsukada, H., Miyasato, K., Nishiyama, S., Fukumoto, D., Kakiuchi, T., & Domino, E. F. (2005). Nicotine normalizes increased prefrontal cortical dopamine D1 receptor binding and decreased working memory performance produced by repeated pretreatment with MK-801: a PET study in conscious monkeys. Neuropsychopharmacology, 30(12), 2144–2153. doi:10.1038/sj.npp.1300745

Tsukada, H., Nishiyama, S., Fukumoto, D., Sato, K., Kakiuchi, T., & Domino, E. F. (2005). Chronic NMDA antagonism impairs working memory, decreases extracellular dopamine, and increases D1 receptor binding in prefrontal cortex of conscious monkeys. Neuropsychopharmacology, 30(10), 1861–1869. doi:10.1038/sj.npp.1300732

Virdee, K., Kentrop, J., Jupp, B., Venus, B., Hensman, D., McArthur, S., … Dalley, J. W. (2016). Counteractive effects of antenatal glucocorticoid treatment on D1 receptor modulation of spatial working memory. Psychopharmacology (Berl), 233(21-22), 3751–3761. doi:10.1007/s00213-016-4405-8

Walsh, T. J., Herzog, C. D., Gandhi, C., Stackman, R. W., & Wiley, R. G. (1996). Injection of IgG 192-saporin into the medial septum produces cholinergic hypofunction and dose-dependent working memory deficits. Brain Res, 726(1-2), 69–79.

Wang, D. C., Lin, H. T., Lee, Y. J., Yu, H. F., Wu, S. R., & Qamar, M. U. (2020). Recovery of BDNF and CB1R in the Prefrontal Cortex Underlying Improvement of Working Memory in Prenatal DEHP-Exposed Male Rats after Aerobic Exercise. Int J Mol Sci, 21(11). doi:10.3390/ijms21113867

Watt, M. J., Roberts, C. L., Scholl, J. L., Meyer, D. L., Miiller, L. C., Barr, J. L., … Forster, G. L. (2014). Decreased prefrontal cortex dopamine activity following adolescent social defeat in male rats: role of dopamine D2 receptors. Psychopharmacology (Berl), 231(8), 1627–1636. doi:10.1007/s00213-013-3353-9

Willig, F., M’Harzi, M., Bardelay, C., Viet, D., & Delacour, J. (1991). Roman strains as a psychogenetic model for the study of working memory: Behavioral and biochemical data. Pharmacology Biochemistry and Behavior, 40(1), 7–16. Retrieved from https://www.embase.com/search/results?subaction=viewrecord&id=L21324338&from=export

Willig, F., Van de Velde, D., Laurent, J., M’Harzi, M., & Delacour, J. (1992). The Roman strains of rats as a psychogenetic tool for pharmacological investigation of working memory: example with RU 41656. Psychopharmacology (Berl), 107(2-3), 415–424. doi:10.1007/bf02245169

Winterfeld, K. T., Teuchert-Noodt, G., & Dawirs, R. R. (1998). Social environment alters both ontogeny of dopamine innervation of the medial prefrontal cortex and maturation of working memory in gerbils (Meriones unguiculatus). J Neurosci Res, 52(2), 201–209. doi:10.1002/(sici)1097-4547(19980415)52:2<201::Aid-jnr8>3.0.Co;2-e

Wojtas, A., Herian, M., Skawski, M., Sobocińska, M., González-Marín, A., Noworyta-Sokołowska, K., & Gołembiowska, K. (2021). Neurochemical and Behavioral Effects of a New Hallucinogenic Compound 25B-NBOMe in Rats. Neurotox Res, 39(2), 305–326. doi:10.1007/s12640-020-00297-8

Yahr, M. D., Duvoisin, R. C., Schear, M. J., Barrett, R. E., & Hoehn, M. M. (1969). Treatment of parkinsonism with levodopa. Arch Neurol, 21(4), 343–354. doi:10.1001/archneur.1969.00480160015001

Zahrt, J., Taylor, J. R., Mathew, R. G., & Arnsten, A. F. (1997). Supranormal stimulation of D1 dopamine receptors in the rodent prefrontal cortex impairs spatial working memory performance. J Neurosci, 17(21), 8528–8535. Retrieved from https://www.ncbi.nlm.nih.gov/pubmed/9334425

Zhang, Q., Jung, D., Larson, T., Kim, Y., & Narayanan, N. S. (2019). Scopolamine and Medial Frontal Stimulus-Processing during Interval Timing. Neuroscience, 414, 219–227. doi:10.1016/j.neuroscience.2019.07.004

